# Periodic actin structures in neuronal axons are required to maintain microtubules

**DOI:** 10.1101/049379

**Authors:** Yue Qu, Ines Hahn, Stephen Webb, Simon P. Pearce, Andreas Prokop

## Abstract

Axons are the cable-like neuronal processes wiring the nervous system. They contain parallel bundles of microtubules as structural backbones, surrounded by regularly-spaced actin rings termed the periodic membrane skeleton (PMS). Despite being an evolutionarily-conserved, ubiquitous, highly-ordered feature of axons, the function of PMS is unknown. Here we studied PMS abundance, organisation and function, combining versatile *Drosophila* genetics with super-resolution microscopy and various functional readouts. Analyses with 11 different actin regulators and 3 actin-targeting drugs suggest PMS to contain short actin filaments which are depolymerisation resistant and sensitive to spectrin, adducin and nucleator deficiency - consistent with microscopy-derived models proposing PMS as specialised cortical actin. Upon actin removal we observed gaps in microtubule bundles, reduced microtubule polymerisation and reduced axon numbers suggesting a role of PMS in microtubule organisation. These effects become strongly enhanced when carried out in neurons lacking the microtubule-stabilising protein Short stop (Shot). Combining the aforementioned actin manipulations with Shot deficiency revealed a close correlation between PMS abundance and microtubule regulation, consistent with a model in which PMS-dependent microtubule polymerisation contributes to their maintenance in axons. We discuss potential implications of this novel PMS function along axon shafts for axon maintenance and regeneration.

**Significance statement:** Axons are cable-like neuronal processes that are up to a meter long in humans. These delicate structures often need to be maintained for an organism’s lifetime, i.e. up to a century in humans. Unsurprisingly, we gradually lose about 50% of axons as we age. Bundles of microtubules form the structural backbones and highways for life-sustaining transport within axons, and maintenance of these bundles is essential for axonal longevity. However, the mechanisms which actively maintain axonal microtubules are poorly understood. Here we identify cortical actin as an important factor maintaining microtubule polymerisation in axons. This finding provides potential explanations for the previously identified, but unexplained, links between mutations in genes encoding cortical actin regulators and neurodegeneration.

## Introduction

Axons are slender, cable-like extensions of neurons which wire the nervous system and propagate nerve impulses (Prokop, 2013). Their damage causes impairment of movement or cognitive abilities (Smith *et al.*, 2000), yet most axons cannot be replaced and their delicate structure usually has to be maintained for an organism’s lifetime. Unsurprisingly, we gradually lose half our axons during healthy ageing and far more in neurodegenerative diseases (Adalbert and Coleman, 2012).

We want to understand the mechanisms of long-term axon maintenance, which will also be relevant for our understanding of axon pathology during ageing and disease. For this, we focus on the cytoskeleton and its immediate regulators, which have prominent hereditary links to neurodegenerative disorders (Prokop *et al.*, 2013). Of particular importance are parallel bundles of MTs which run the length of axons; they need to be actively maintained because they form the structural backbones of axons and highways for life-sustaining transport of materials, organelles and signals between cell bodies and the distal synaptic endings (Prokop, 2013; Voelzmann *et al.*, 2016b).

F-actin is a potent regulator of MTs in neuronal contexts, such as in growth cones or axonal branching (Prokop *et al.*, 2013; Kalil and Dent, 2014), but it is unknown whether F-actin might also be a regulator during MT maintenance in axons. Most of the actin networks occurring in axons appear ill-suited for such a task because they are locally restricted or short-lived, including dense networks at the axon initial segment (Rasband, 2010; Li *et al.*, 2011; Galiano et al., 2012; Watanabe *et al.*, 2012), shaft filopodia (Kalil and Dent, 2014), lamellipodia-like actin waves (Flynn *et al.*, 2009), or transiently occurring longitudinal actin filaments *(Ganguly et al.*, 2015). The only known persistent actin networks occurring all along axons is the recently-discovered periodic membrane skeleton (PMS), which can be observed in culture and *in vivo*, in different animal species, but also in dendrites or neurite-like glial processes (Xu *et al.*, 2013; Lukinavičius *et al.*, 2014; D’Este *et al.*, 2015; He *et al.*, 2016).

PMS is believed to represent a specific form of cortical F-actin: it consists of short, adducin-capped actin filaments which are bundled into rings and cross-linked by spectrins that space them into regular ~180 nm intervals. However, this model is mainly based on super-resolution microscopical analyses, and very few actin regulators have been functionally assessed so far for their potential contributions to PMS architecture. Thus, it was suggested that spectrins play a major role in their formation (Zhong *et al.*, 2014), whereas knock-down of ankyrinB (Zhong *et al.*, 2014) or loss of adducin (Leite *et al.*, 2016) had no effect on PMS organisation.

Here we took a new approach to analyse PMS. Analyses performed so far made use of STORM or STED microscopy, whereas we used structured illumination microscopy (SIM). SIM provides slightly lower resolution but gives highly robust readouts for the periodic patterns which enabled us to perform quantitative analyses of PMS abundance across axon populations. As our cellular system we used neurons of the fruit fly *Drosophila* which provide easy access to experimental and genetic analyses (Prokop *et al.*, 2013), and so we were able to study the functional depletion of 11 actin regulators and 3 actin-targeting drugs. We found a range of robust and highly reproducible effects on PMS which provide functional support for the view that PMS represents cortical actin specialisations. We then combined these actin manipulations with a number of readouts for axonal MTs, suggesting prominent roles for cortical actin networks in promoting the polymerisation of axonal MTs relevant for axon maintenance.

## Materials & Methods

### Fly stocks

All mutant alleles used in this study are well characterised. The following loss-of-function mutant alleles were used:

The *Hu li tai* shao/adducin loss-of-function mutant allele ***hts***^1^ and ***hts***^01103^ are both strong hypomorphic alleles due to a transposable element insertion. They were isolated in two independent single P-element mutagenesis screen (Yue and Spradling, 1992; Spradling *et al.*, 1999). Our experiments studying filopodial length were carried out with 6 days pre-culture to deplete maternal contribution (Sánchez-Soriano *et al.*, 2010).

The *α-pectrin* allele *α-**Spec**^rg41^* is a protein null allele caused by a 20 bp deletion, resulting in a premature amber stop codon near the 5’ end of the coding region (Lee *et al.*, 1993). Our staining with anti-á-Spectrin antibody failed to detect any signal in *α-**Spec**^rg41^* mutant primary neurons (data not shown) (Hülsmeier *et al.*, 2007).

The *β-Spectrin* allele *β-**Spec***^S012^ is a protein null allele caused by nucleotide substitution C538T with reference to the 2291aa isoform (Hülsmeier *et al.*, 2007). Hemizygous *β-Spec*^S012^ mutant embryos lack detectable β-Spectrin expression (Hülsmeier *et al.*, 2007). Our experiments studying filopodial length were carried out with 6 days pre-culture to deplete maternal contribution (Sánchez-Soriano *et al.*, 2010).

The *ankyrin2* null mutant allele ***ank2**^518^* is a transposon insertion in the 6^th^ intron of the *ank2* gene causing the disruption of all three Ank2 isoforms (Pielage *et al.*, 2008).

SCAR (homologue of human WASF1-3) and HEM-protein/Kette (homologue of human NCKAP1/NAP1) are essential components of the WAVE/SCAR complex required for Arp2/3-mediated nucleation in *Drosophila* neurons (Schenck *et al.*, 2004). The mutant allele ***Hem***^03335^ is a protein null caused by a P-element insertion 39bp downstream of the putative transcription start site (Baumgartner *et al.*, 1995; Schenck *et al.*, 2004). The ***SCAR**^Ä37^* deletion is a protein null allele caused by imprecise P-element excision (Zallen *et al.*, 2002; Schenck *et al.*, 2004).

Arpc1/Sop2 is the homologue of the essential regulatory Arp2/3 subunit ARPC1B/p41. The mutant ***Arpc1***^1^ (= *Sop2*^1^; from B. Baum) allele is caused by a 207bp genomic deletion that removes the last 62 codons of *arpc1* (Hudson and Cooley, 2002).

***DAAM**^Ex68^* is a null allele generated via imprecise P-element excision resulting in deletion of the C-terminal 457 amino acids, including sequences corresponding to the ‘DAD’ domain and most of the ‘FH2’ domain (Matusek *et al.*, 2006).

The *enabled* mutant allele ***ena**^23^* is caused by a nucleotide exchange introducing a STOP codon leading to a 52aa C-terminal truncation that deletes the EVH2 domain required for tetramerisation of Ena (Ahern-Djamali *et al.*, 1998). In *ena*^23^ mutant embryos, anti-Ena staining (clone 5G2, mouse) is strongly reduced in primary neurons, CNSs and tendon cells (Alves-Silva *et al.*, 2008; Sánchez-Soriano *et al.*, 2010; Gonçalves-Pimentel *et al.*, 2011).

The *chickadee/*profilin mutant null allele ***chic***^221^ is caused by an intragenic deletion removing 5’ non-coding and some of coding region of *chic* (Verheyen and Cooley, 1994; Wills *et al.*, 1999); anti-Chic staining (mouse, clone chi1J) is strongly reduced in *chic*^221^ mutant CNS and primary neurons (Gonçalves-Pimentel *et al.*, 2011).

The two chemically induced *short stop* mutant alleles ***shot**^3^* and ***shot**^sf20^* are widely used and are the strongest available (likely null) alleles (Kolodziej *et al.*, 1995; Prokop *et al.*, 1998).

For live imaging with the ***UAS-eb1-GFP*** line (courtesy of P. Kolodziej) (Sánchez-Soriano *et al.*, 2010) we used the pan-neuronal driver line ***sca-Gal4*** driver line for experiments at 6-8HIV (Sánchez-Soriano *et al.*, 2010), and for experiments >1DIV we used ***elav-Gal4*** (3^rd^ chromosome) driver lines (Luo *et al.*, 1994).

Green balancers used were *Kr::GFP* (Casso *et al.*, 2000) and *twi::GFP* (Halfon *et al.*, 2002).

### Cell culture

Primary neuron cultures were generated following procedures that were described in detail in previous papers (Sánchez-Soriano *et al.*, 2010; Prokop *et al.*, 2012; Beaven *et al.*, 2015). In brief, embryos were dechlorinated using bleach, selected for the correct genotypes at about stage 11 using fluorescent balancer chromosomes (stages according to Campos-Ortega and Hartenstein) (Campos-Ortega and Hartenstein, 1997), sterilised with ethanol, and mechanically crushed. Resulting cells were chemically dispersed and then washed in Schneider’s medium. They were either directly plated, or kept in centrifuge tubes for 3-7 days before plating in order to deplete maternal protein product (pre-culture). In both cases, cells were plated at standard concentration onto glass coverslips which were either uncoated or coated with Concanavalin A. Coverslips were kept on special incubation chambers where cells were grown as hanging drop cultures at 26°C. For analyses at the growth cone stage, cells were grown for 6-8HIV (hours *in vitro)* on glass or ConA which were extended to 20HIV for pre-cultured neurons (always on ConA). Mature neurons were analysed at 3-10 days (always on ConA). To deplete maternal gene product, cells were pre-cultured in Schneider’s medium in centrifuge tubes for up to 7 days before they were plated out as described above.

For drug treatments, solutions were prepared in a cell culture medium from stock solutions in DMSO. Cells were treated with 200nM Latrunculin A (Biomol International) for 1HIV or 4HIV, 800nM or 1.6 ìM Cytochalasin D (Sigma) for 1HIV or 4HIV, 20 iM nocodazole (Sigma) for 2.5HIV, or 10μM SMIFH2 (Sigma) for 4HIV, respectively. For controls, equivalent concentrations of DMSO were diluted in Schneider’s medium.

### Immunohistochemistry

Primary fly neurons were fixed in 4% paraformaldehyde (PFA) in 0.1 M PBS (pH 6.8 or 7.2) for 30 min at room temperature (RT), then washed three times in PBS with 0.3% TritonX-100 (PBT), followed by staining.

Antibody and actin staining and washes were performed in PBT using anti-tubulin (clone DM1A, mouse, 1:1000, Sigma; alternatively, clone YL1/2, rat, 1:500, Millipore Bioscience Research Reagents), anti-α-spectrin (clone 3A9, mouse, 1:200, DSHB), anti-elav (clone 7E8A10, rat, 1:1000, DSHB), anti-Synaptotagmin; FITC-, Cy3- or Cy5-conjugated secondary antibodies (donkey, purified, 1:200; Jackson Immuno Research), TRITC/Alexa647-, FITC-conjugated phalloidin (1:200; Invitrogen and Sigma). Specimens were embedded in Vectashield.

For stimulated emission depletion (STED) and structured illumination microscopy (SIM), cells were cultured for 8HIV or up to 10DIV (days in vitro) at 26°C on ConA-coated 35 mm glass-bottom MatTek dishes (P35G-0.170-14-C). Cells were fixed with 4% PFA, washed 3 times in PBT, then stored for transport in PBS sealed with Parafilm. Before imaging, cells were incubated for 1hr with 2iM SiR-actin in PBS (Spirochrome) (Lukinavičius *et al.*, 2014), then washed once with PBS.

### Microscopy

SIM was performed with an Elyra PS1 microscope (Zeiss) with a 100× oil immersion objective lens (NA = 1.46) and 642nm diode laser. Raw images were acquired with three or five grating angles.

STED was performed using a Leica SP8 gated STED microscope with a 100× oil immersion objective lens (NA = 1.40), 640 nm excitation and 775 nm depletion.

Standard images were taken with a 100×/1.30 oil iris Ph3 8/0.17 objective (plus 1.6× Optovar) on an Olympus BX50WI microscope with filter sets suitable for FITC, Cy3 and Cy5 and equipped with a AxioCam camera (Zeiss). 2012 software (blue edition, Zeiss) was used to acquire images.

Time-lapse live imaging of Eb1::GFP-expressing cultured neurons was performed on standard ConA-coated cover slips under temperature-controlled conditions (26°C) on a Delta Vision RT (Applied Precision) restoration microscope with a 100×/1.3 Ph3 Uplan Fl phase objective, Sedat filter set (Chroma 89000), and a Coolsnap HQ (Photometrics) camera. Images were taken every 4s for 2-3mins with exposure times of 0.5-1s, and were constructed to movies automatically.

To observe the impact of CytoD treatment on axon retraction, Eb1::GFP expressing primary neurons were cultured on 35mm glass-bottom MatTek dishes. About 10 cells per slide were filmed one-by-one for 2 mins before 800nM or 1.6 ìM CytoD was applied, and then revisited for further imaging at 0hr, 0.5HIV, 1hr, 1.5HIV and 2HIV after application.

### Data analysis

The relative abundance of periodic axonal actin structures (PMS) was assessed on randomly chosen SIM images containing axons of SiR-actin-stained primary neuronal cultures, achieving sample numbers usually above 300. These were taken from 4 independent culture preparations obtained from at least 2 independent experimental repeats performed on different days. From each single culture preparation a minimum of 20 SIM images was obtained, and in each image all neurite segments of >6μm length were counted. Generally, these neurite segments showed a consistent presence or absence of PMSs all along. To avoid bias, image analyses were performed blindly, i.e. the genotype or treatment of specimens was masked.

We can be certain that analysed neurites represent axons. Thus, dendrites in our culture system are located on neuronal cell bodies but develop sparsely when analysed at 3DIV (Sánchez-Soriano *et al.*, 2005). We found the same for cultures at 10 DIV where only 15% of somata display short processes (5.8+/-0.8μM) which lack the presynaptic marker Synaptotagmin typically found only in axons. Therefore, by choosing neurites of >6μm length located at sufficient distance from cell bodies, we are confident that all images represent axonal segments.

Filopodial lengths were measured using the segmented line tool in ImageJ. We included filopodia along axon shafts and at growth cones, but excluded those on cell bodies.

MT disorganisation was assessed as MT disorganisation index (MDI), as first published here: the area of disorganisation was measured using the freehand selection in ImageJ. This value was then divided by axon length (measured using the segmented line tool in ImageJ) multiplied by 0.5ìm (typical axon diameter), i.e. approximating axon area without disorganisation.

**Figure 3.**
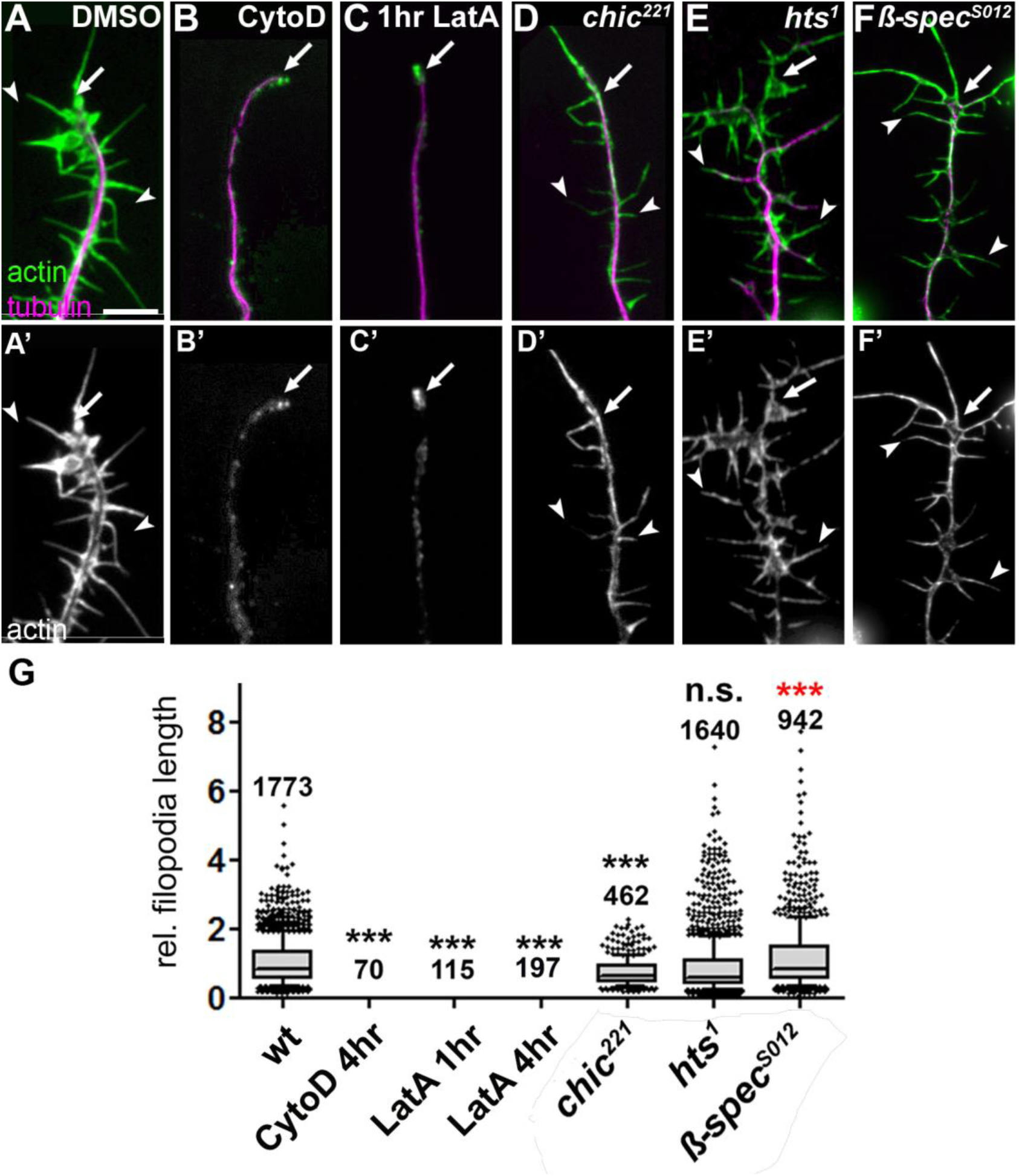
Effects of actin manipulations on filopodial length. (**A-F**’) Filopodial length phenotypes in DMSO-treated wildtype primary neurons, or neurons treated with drugs or being mutant, as indicated; cells are double-labelled for actin (green in top row, white in bottom row) and tubulin (magenta in top row); drug treatments: 800nM CytD for 4hrs, 200nM LatA for 1hr. (**G**) Quantifications of filopodia length caused by drug treatment or mutations shown on left (all normalised and compared to DMSO-treated controls); numbers above the bars indicate the numbers of filopodia analysed in each experiment; note that filopodia were completely absent in all cases of CytoD and LatA treatment. P values were calculated using the Mann-Whitney Rank Sum test (NS: P>0.05, *: P<0.05, **: P<0.01,***: P<0.001), red indicate higher than wildtype). Scale bar in A represents 10μm in A-F.

To assess frequencies of neurons with axons (Fig. 3), we double-labelled primary neurons for tubulin and the neuron-specific nuclear protein Elav (Robinow and White, 1991). When defining an axon as a tubulin-stained process longer than the diameter of the soma (Gonçalves-Pimentel *et al.*, 2011), 76% of Elav-positive neurons display an axon in wildtype control cultures.

To assess frequencies of gaps in axons, relative numbers of primary neurons were counted where anti-tubulin along axons was discontinuous, i.e. displaying gaps (Figure 3) (Alves-Silva *et al.*, 2012).

EB1::GFP live analyses upon actin drug treatments were performed as described previously (Alves-Silva *et al.*, 2012). To measure speed, EB1 comets were tracked manually using the “manual tracking” plug-in of ImageJ. To analyse the number of comets, EB1 spots within an axon region of interest were counted over the whole captured time period at each of the different time points (i.e. 5, 30, 60, 90 and 120 mins after drug application), and the means of these comet numbers per axon region and time point were normalised to the mean of comet numbers in this same region before drug treatment. For each time point and treatment 6-14 different axons were analysed. Error bars shown graphs (Figure 6C-H) indicate SEM of normalised data of all axons for each time point.

**Figure 6.**
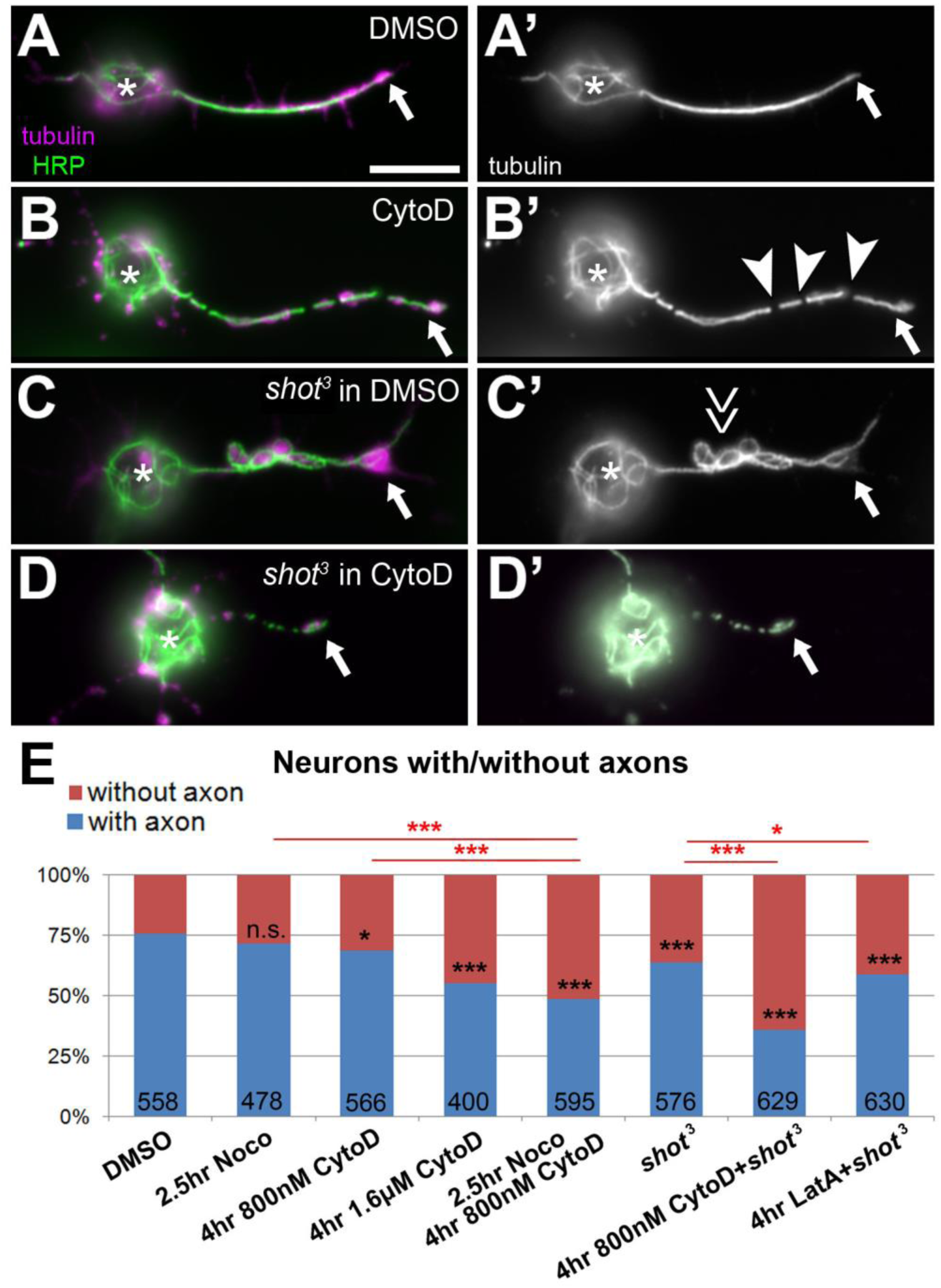
Combined loss of F-actin and Shot reduces axon numbers. **A-D**') Primary neurons at 8HIV from wildtype (wt) or *shot* mutant embryos at 8HIV treated for 4hrs with either DMSO or CytoD as indicated on the left and stained for tubulin (green, white on right) and HRP (magenta). CytoD treatment alone causes gaps in the axonal tubulin staining (B; arrowheads, MT gaps), Shot deficiency causes MT disorganisation (C; double arrow, MT disorganisation); combination of *shot* with CytoD has a detrimental effect on axons (D). (**E**) Statistical analysis of neurons with/without axons; numbers in bars refer to analysed neurons; all data were compared to DMSO controls via X2 analysis (NS P>0.050, *P<0.050, **P<0.010,***P<0.001; for detailed data see Tab.S1). Scale bar in A represents 10μm in A-D.

For all analyses, GraphPad Prism 5 was used to calculate the mean and standard errors of the mean (SEM), and to perform statistical tests using either the Mann-Whitney U test or the Chi^2^ test.

**Figure 1.**
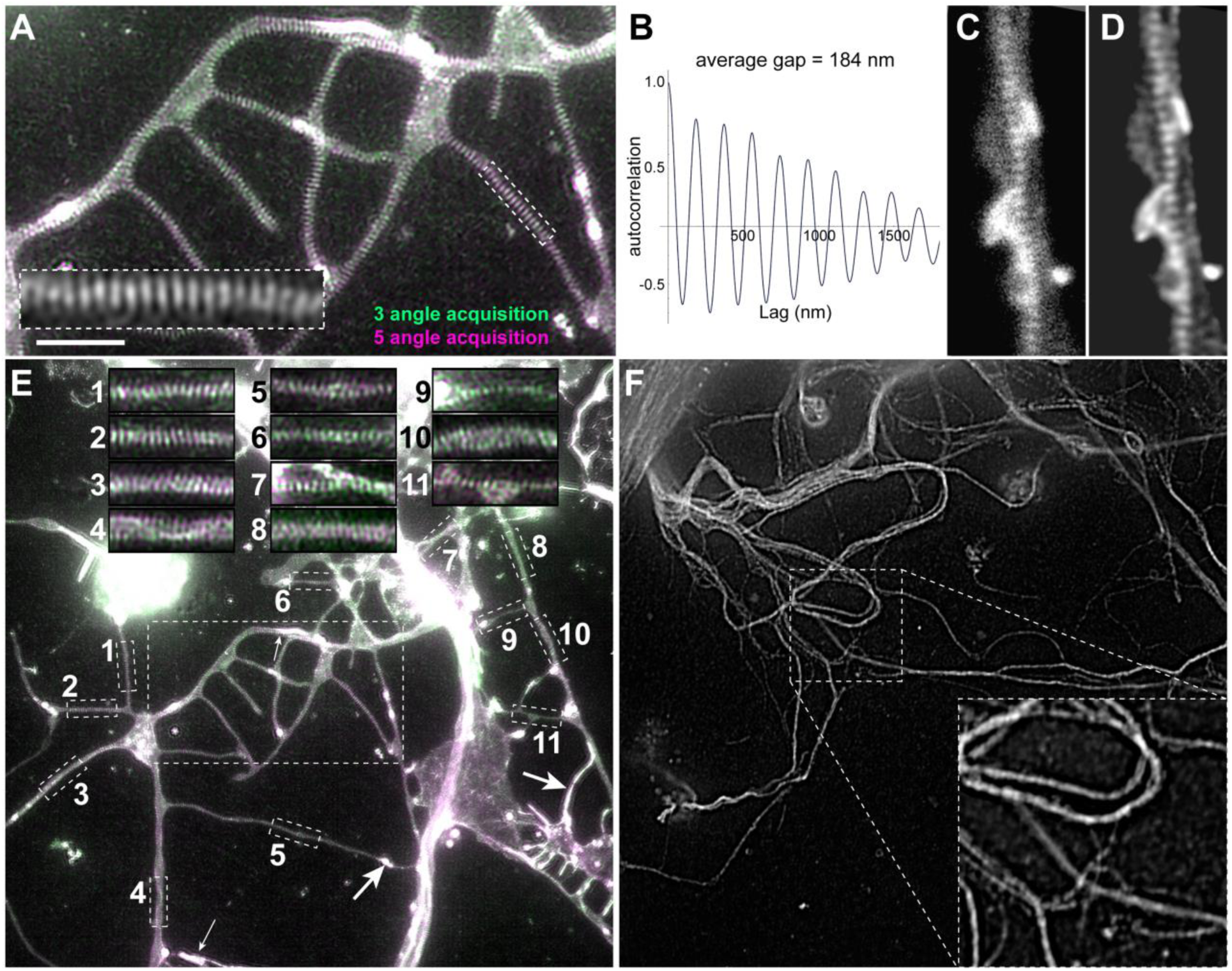
Super-resolution images of *Drosophila* axons display PMS. (**A**) SIM image of SiR-actin-stained *Drosophila* primary neurons at 10DIV; the image shows precise overlay of two independent rounds of image acquisition using three (green) versus five (magenta) rotation angles; the framed area is shown 4-fold magnified in the inset. (**B**) Autocorrelation analysis showing the regular periodicity of the actin staining with a lag of 184nm. (**C**,**D**) Axon visualised via STED shown as raw (C) and deconvolved image (D). (**E**) Full SIM image of SiR-actin-stained neurons at 10DIV; the large emboxed area is shown at larger scale in A; small emboxed areas are shown as insets at the top illustrating the high reliability of PMS appearance in these cultures; arrows mark dotted or elongated actin accumulations and emboxed area 4 might show an actin trail. (**F**) Full SIM image of neurons at 10DIV stained with anti-Tubulin; emboxed area is shown as 2.4 fold magnified inset at bottom right (note that tubulin staining does not show any periodicity). Scale bar in A represents 3ìm in A and the inset of F, 1.2μm in C, D and inset of A, 7.2μm in E and F, and 2.1μm in insets of E.

To generate the average autocorrelation curves (Figs. 1E), intensity profiles were extracted from typical axon images using Image J, then analysed using Mathematica 10.2. The autocorrelation function *R*(*τ*,**x**_*i*_) at a lag of *τ* for an equally spaced *n*_*i*_ point data series, x_*i*_ = 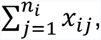 is given by,

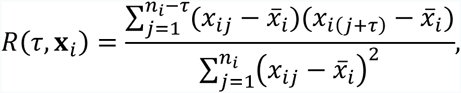

where 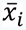 is the mean of **x**_*i*_. For each profile the autocorrelation function is calculated, these are then weighted by the length of the profile to give an average autocorrelation function,

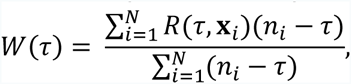

where *N* is the total number of profiles being averaged over.

To compare the intensity of autocorrelation curves between different conditions, we performed Mann-Whitney tests on the autocorrelation magnitudes at each point, and then applied Fisher’s Method with correlation (Dai *et al.*, 2014) to calculate an overall p-value.

## Results

### *Drosophila* periodic membrane skeleton in axons is a gradually developing, stable actin network

To visualise axonal actin, we used structured illumination microscopy (SIM) and imaged primary neurons of the fruit fly *Drosophila* which were cultured for > 6DIV (days *in vitro)*, then fixed and stained with SiR-actin. We used strategies which allowed us to selectively visualise axons and exclude dendrites (see Methods). These SIM-imaged axons revealed irregular dotted or elongated actin accumulations potentially demarcating synapses, occasional longitudinal actin trails as described also for mammalian neurons (emboxed area 4 in Fig. 1E) (*Ganguly et al., 2015)*, and abundant periodic actin patterns with a repeat length of 184±2nm, highly reminiscent of the periodic membrane skeleton (PMS; Fig. 1A-E). Further validation clearly showed the PMS to be genuine. Firstly, we found the same periodicity when using stimulated emission depletion (STED) microscopy (Fig. 1C,D). Secondly, SIM imaging with anti-tubulin staining showed no periodicity (Fig. 1F). Thirdly, overlay images of the same preparation taken from three versus five angles showed a clear overlay (Fig. 1A,E). Finally, our findings in cultured primary neurons are in agreement with *in vivo* observation of PMS in axons of fly embryos and larvae (He *et al.*, 2016) and periodic spectrin patterns observed in earlier studies (*Pielage et al.*, 2008).

Compared to STED, SIM provides slightly lower resolution and does not permit measurements of actin content in PMS, since it uses a demodulation algorithm with which the raw detected signal cannot be converted into photon counts. However, as an essential advantage for our studies, SIM allowed fast imaging and revealed PMS with high reliability in virtually all preparations (see overview image Fig. 1E). This enabled us to reach sample numbers of several hundred to over a thousand axon segments from different biological and technical repeats of each set of experiments. These conditions were ideal for systematic quantitative analyses, and we quantified the relative number of axon segments displaying PMS (termed “PMS abundance”) across axon populations of each experimental condition (see Methods).

**Figure 2.**
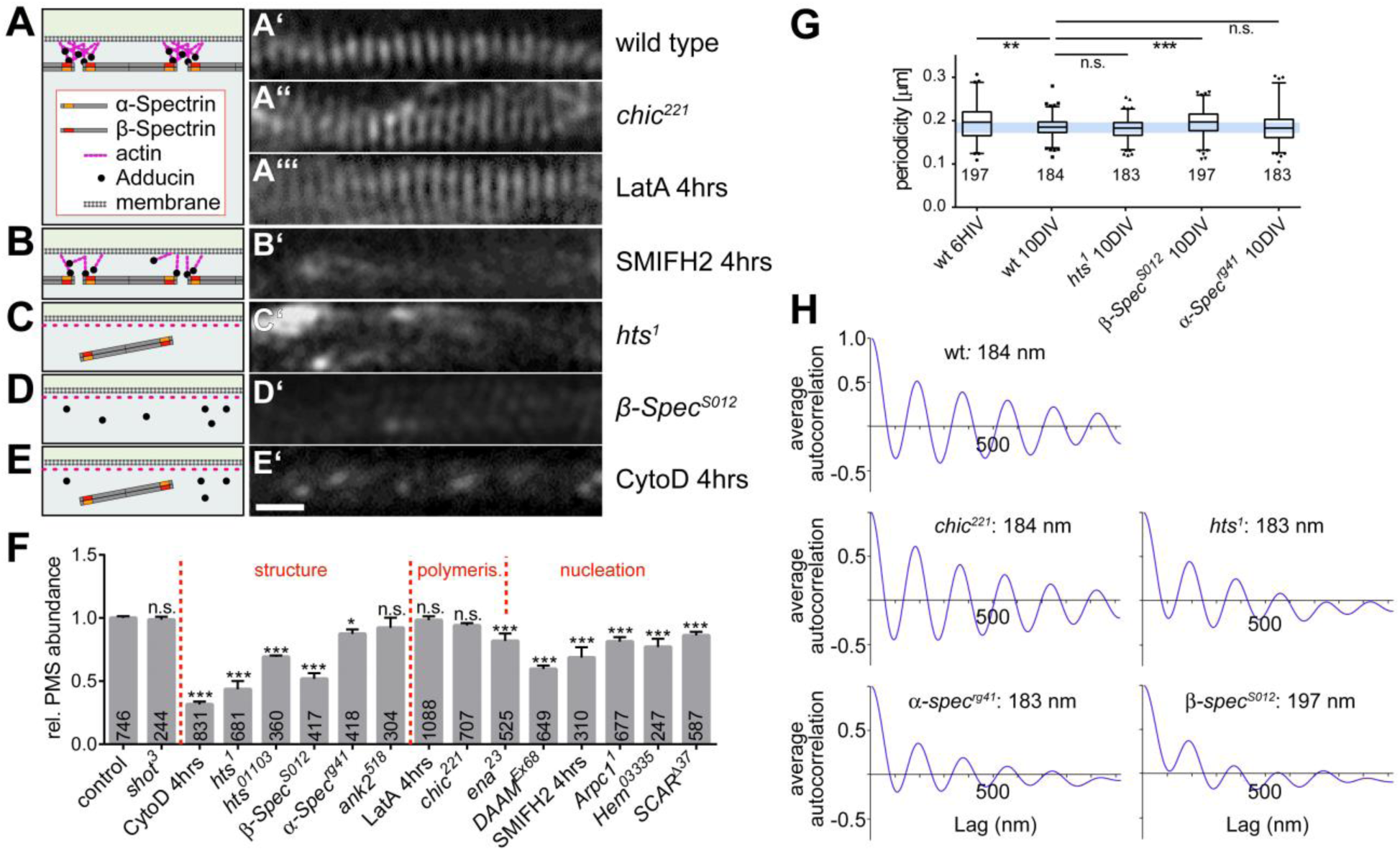
Functional dissection of PMS. (**A-E**) Representative SIM images of SiR-actin-labelled axons at 10DIV genetically or pharmacologically manipulated as indicated on the right (scale bar in E’=550nm for all SIM images); schematics on the left provide an interpretation of the observed phenotype, based on the previously proposed cortical actin model (Xu *et al.*, 2013). (**F**) Quantification of PMS abundance in axons of mature neurons at 10DIV, sorted by manipulations affecting structure, polymerisation or nucleation; P values were obtained via X^2^ analysis of raw data comparing axon segments with/without PMS (NS: P>0.05,*: P=0.05; **: P=0.01; ***: P=0.001); numbers in bars represent sample numbers (i.e. analysed axon segments); error bars represent SEM of independent experimental repeats. (**G**) Box and whisker plot showing periodicity within MPS (whiskers indicate 90^th^ percentile); note that the lower whiskers are truncated by the limitation of image resolution achievable in SIM; P values were obtained via X2 analysis of raw data comparing axon segments with/without PMS (NS: P>0.05,*: P=0.05; **: P=0.01; ***: P=0.001). (H) Average spatial autocorrelation curves showing the PMS periodicity, each calculated from 15 axon segments showing clear PMS. Detailed data underlying graphs of this figure are given in Tab.S1.

Using PMS abundance as readout, we investigated the relative vulnerability/resistance of PMS to the application of actin-destabilising drugs. For mammalian axons it was shown that applications of the actin-destabilising drugs cytochalasin D (CytoD) and latrunculin A (LatA) at high doses destroy PMS (Zhong *et al.*, 2014). Of these two drugs, CytoD is known to sequester actin monomers and also to directly destabilise barbed ends of actin filaments (Peterson and Mitchison, 2002). Accordingly, when we treated neurons at 10DIV with a concentration as low as 800nm, we found a significant reduction of PMS abundance down to 32% as compared to wild type (Fig. 2E’, F).

In contrast, LatA only sequesters actin monomers, thus primarily suppressing their polymerisation (Peterson and Mitchison, 2002). Accordingly, when we used 4hr low-dose treatments with LatA at 200nM, this caused only a non-significant reduction to 94% (Fig. 2A’’’,F). As a positive control, we used young neurons which, in control cultures, display prominent filopodia dependent on highly dynamic actin filament networks (Gonçalves-Pimentel *et al.*, 2011). We found that 4hr and even 1hr treatments with either LatA or CytoD completely eliminated filopodia (Fig. 3B,C,G), clearly demonstrating that both drugs were highly effective at the low dosage used.

The differential effects that especially LatA had on dynamic actin networks in filopodia, but not on the structure of PMS, suggests that the barbed ends of actin filaments in PMS are to a degree protected against actin monomer sequestration. A candidate protein mediating this effect could be adducin (see next section).

### PMS abundance strongly depends on the cortical actin regulators spectrin and adducin

We next used SIM analyses of neurons at 10DIV and studied known components of cortical actin, in particular spectrins, adducin and ankyrin (Baines, 2010), which were all reported to reside in or at PMS (Xu *et al.*, 2013; Zhong *et al.*, 2014).

We first tested spectrins which usually form hetero-oligomers composed of α- and ß-spectrins. Whilst there seems to be no reported data for α-spectrin in mammalian neurons, ß-spectrin was shown to localise at PMS, and it has been stated that knock-down of ßll-spectrin eliminates PMS (Xu *et al.*, 2013; Zhong *et al.*, 2014). *Drosophila* α- and ß-Spectrin are each encoded by only one gene, and both proteins are localised in axons (Pielage *et al.*, 2006; Garbe *et al.*, 2007; Hülsmeier *et al.*, 2007). We therefore tested neurons carrying the *α-Spec^rg41^* or *β-Spec*^*S012*^ null mutant alleles which eliminate *Drosophila α-* and ß-Spectrin, respectively (see Methods). We found *β-Spec*^*S012*^ to cause a drastic and highly significant reduction of PMS abundance to 58% (Fig. 2D’,F). Furthermore, within existing PMS, the average periodic spacing was highly significantly increased to 197±2 nm and it was less regular: autocorrelation curves averaged from 15 independent samples (see Methods) indicated strongly reduced periodicity (Figs. 2G,H and S1). In contrast, *α*-Spec^*rg41*^ causes only a moderately significant and far milder reduction of PMS abundance to 89% (Fig. F), and average periodicity of 183±3 nm was normal. However, there was a larger variation in distances as was also reflected in the lower amplitude in the averaged autocorrelation curve (Figs. 2G,H and S1). These findings suggest that α/ß-spectrin heter-oligomers contribute to PMS abundance and organisation, with ß-spectrin playing a more important role as judged from the stronger phenotypes.

Our findings are in agreement with other reports which used very different readouts to conclude that α-spectrin is of less importance in *Drosophila* neurons than ß-spectrin; ß-spectrin maintains a good degree of functionality even in the absence of alpha (Pielage *et al.*, 2006; Garbe *et al.*, 2007; Hülsmeier *et al.*, 2007). Mechanistically, this can be explained through the fact that ß-spectrins, but not μ-spectrins, contain the essential binding sites for actin and adducin, and ß-spectrin at vertebrate synapses can form functional homo-oligomers (Bloch and Morrow, 1989; Pumplin, 1995).

Adducin caps F-actin barbed ends and is therefore a likely candidate for providing a mechanism that renders PMS less sensitive to CytoD and LatA. Surprisingly, complete loss of mammalian adducin function was reported neither to affect the periodicity of PMS nor the expression of spectrin (Leite *et al.*, 2016). However, this study did not investigate the abundance of PMS. We therefore analysed PMS in neurons carrying the *hts*^1^ null mutant allele which abolishes *Drosophila* adducin. As observed in mammalian axons, we found that PMS in these neurons had a normal periodicity of 183±2 nm (Fig. 2G, H) with a moderate deviation in reliability (Fig. S1), and axonal α-spectrin staining seemed unaffected in *hts*^1^ mutant neurons (Fig. S2). However, in contrast to the normal appearance of these qualitative measures, our quantitative analyses of *hts*^1^ mutant neurons revealed a strong reduction of PMS abundance to 43.5% (Fig. 2C’, F). To confirm this finding, we used the independent strong loss-of-function mutant allele *hts*^01103^ and likewise found a highly significant reduction in PMS abundance to 69.2% (Fig. 2F).

Finally, we tested ankyrin which is known to link cortical actin to the membrane and other structures, but was reported to not affect PMS formation in mammalian axons (Zhong *et al.*,2014). Similarly, our quantitative analyses using the null mutant *ank2*^*518*^ allele showed only a mild, non-significant reduction in PMS abundance to 92% (Fig. 2F).

Taken together, two key players of cortical F-actin networks, spectrin and adducin, are important for PMS formation and/or maintenance. Notably, the *hts*^1^ and *β-Spec*^S012^ mutant alleles affect PMS but cause no reduction in filopodial length in young neurons (maternal contribution removed; Fig. 3E,F,G). This, together with our pharmacological studies, lends further support to the notion that actin networks of PMS significantly differ from dynamic actin networks.

### Actin filaments in PMS are likely short but high in number

Our 4hr treatments with the actin monomer sequestering drug LatA already suggested that PMS is less dependent on actin polymerisation than other actin networks. This is consistent with PMS actin filaments being relatively short, as proposed previously (Xu *et al.*, 2013). They should therefore have little requirement for actin elongation factors. We tested this by analysing neurons carrying the *chic*^221^ null mutant allele which eliminates the function of *Drosophila* profilin, a prominent actin elongation factor (Gonçalves-Pimentel *et al.*, 2011). As predicted, the *chic*^221^ mutant neurons displayed no significant reduction in PMS abundance at 10DIV (Fig. 2A’’,F), and the autocorrelation intensity has no significant difference from the wildtype (Figs. 2G,H and S1). Next we tested the involvement of Ena/VASP which has a number of functions, one being to closely collaborate with profilin in actin filament elongation (Bear and Gertler, 2009; Gonçalves-Pimentel *et al.*, 2011). However, when using the *ena*^23^ allele to deplete Ena/VASP, we found a significant reduction in PMS abundance at 10DIV to 83% (Fig. 2F). Therefore, further profilin-independent functions of Ena/VASP seem to play a role at PMS. One of these roles is to promote nucleation (the formation of new actin filaments) (Gonçalves-Pimentel *et al.*, 2011), and we reasoned that actin nucleation should be particularly important for PMS where actin filaments are short and therefore expected to be high in number.

To test this, we depleted the functions of two distinct nucleators, Arp2/3 and the formin DAAM, which are known to function in parallel in *Drosophila* primary neurons (Gonçalves-Pimentel *et al.*, 2011). We found that both nucleators contribute to PMS. First, genetic depletion of three proteins required for Arp2/3 function (see Methods) showed a consistent reduction of PMS abundance to ~80% (Fig. 2F; *Arpc1^1^:* 79%; *SCAR^Ä37^:* 85%; *Hem^03335^:* 75%). Second, neurons carrying the *DAAM*^Ex68^ null mutation or treated with the formin-inhibiting drug SMIFH2 (4hrs at 10μM), showed an even stronger reduction to 60% and 68% (Fig. 2B’,F).

Therefore, our data support the notion that PMS contains short actin filaments (Xu *et al.*, 2013), and we propose that these filaments are accordingly high in number and therefore very dependent on actin nucleation. Notably, the alleles used here further suggest that PMS is fundamentally different from other actin networks: the profilin-deficient *chic*^221^ allele which hardly affects PMS (Fig. 3D,G) causes severe shortening of filopodia (Gonçalves-Pimentel *et al.*, 2011), whereas loss of nucleators clearly affects PMS but not the length of filopodia (Gonçalves-Pimentel *et al.*, 2011).

### PMS in axons of young, growing neurons appear less stable

It was reported for mammalian neurons that PMS are gradually established during the first days of development (Xu *et al.*, 2013; D’Este *et al.*, 2015). We therefore studied the developmental timeline and found a gradual increase in PMS abundance from 20% at 6HIV (hours *in vitro)* to 82% at 10DIV (Fig. 4A,C). However, PMS in early neurons displayed a larger average spacing of 197±4 nm, with extreme cases reaching up to 300 nm distance between actin peaks, as is also reflected in an almost complete lack of information in averaged autocorrelation curves for these neurons (data not shown). This is reminiscent of the irregularities observed in *β-Spec*^S012^ mutant neurons at 10DIV (Figure 2G,H) and might indicate structural immaturity.

We therefore studied whether young PMS might deviate from mature PMS in other ways, and challenged them with some of the actin manipulations we used at 10DIV (Fig. 4C). In agreement with studies at 10DIV, we found that neurons at 8HIV displayed no changes in PMS abundance when depleted of profilin (*chic*^221^; 96±0.5%), but a strong reduction in PMS abundance when treated for 4hrs with 800nM CytoD (38±4.4%), when carrying the *DAAM^Ex68^* mutant allele (66±8.6%), or the *ß-Spec^S012^* mutant (66±8.5%). Therefore, PMS in young axons seem to contain many short actin filaments which are in the process of being organised through spectrins. However, in deviation from old neurons, 4hr treatment of young *Drosophila* neurons with 200nM LatA highly significantly reduced PMS abundance to 48%, suggesting that young PMS are more reliant on actin polymerisation. To test whether PMS are nevertheless qualitatively different from other dynamic actin networks, we reduced LatA application to 1hr, which is time enough to effectively eliminate filopodia and actin patches along axon shafts (Fig. 3C), but is insufficient to cause significant PMS loss (Fig. 4B, D).

**Figure 4.**
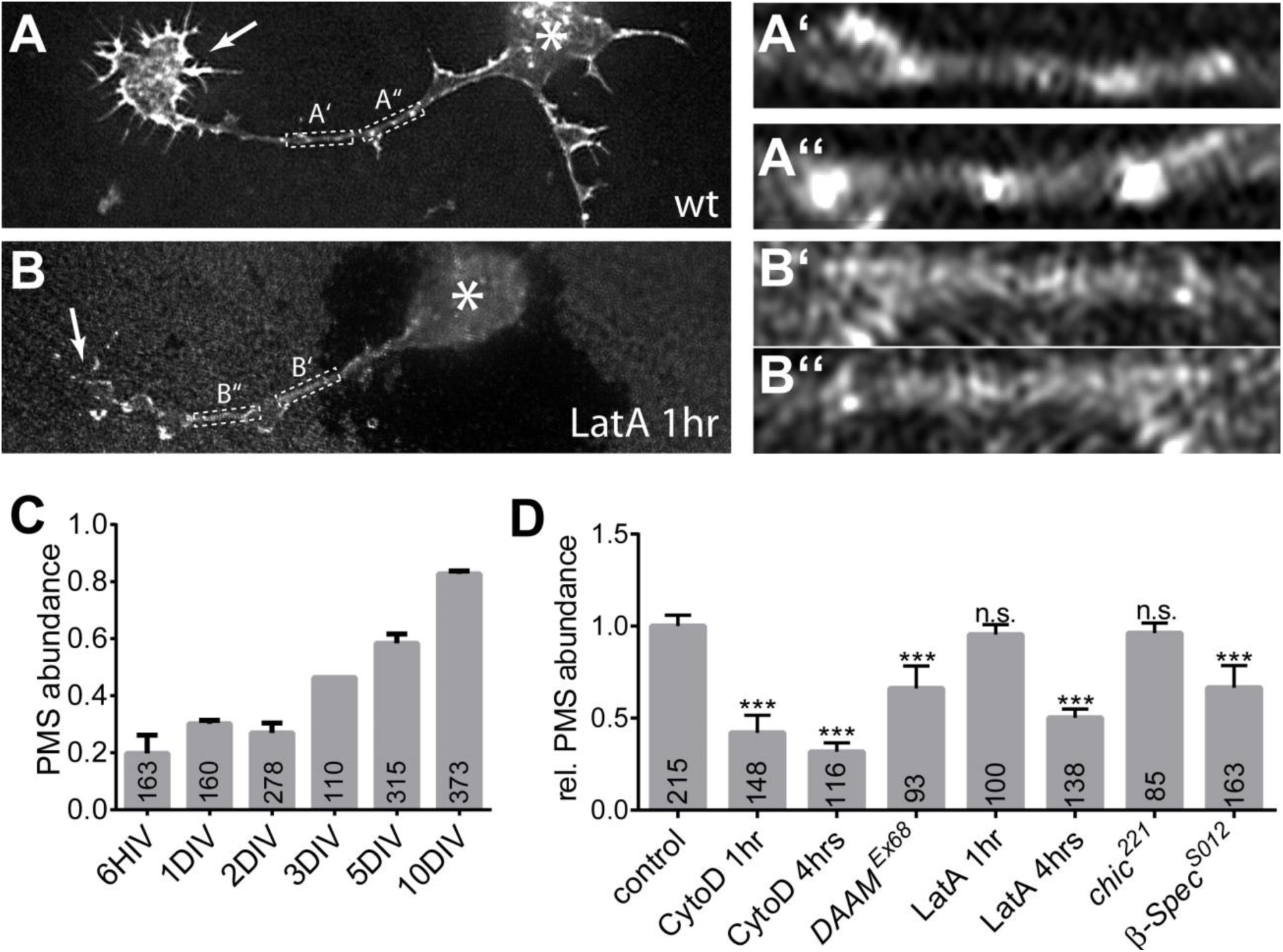
(**A**,**B**) Examples of PMS in axons of neurons at 6HIV, either untreated (A) or treated with 200nM LatA for 1hr (B); asterisks indicate cell bodies, arrowheads the growth cones, framed areas are shown 8-fold enlarged on the right (A’-B”). Note that actin accumulations at growth cones and dotted actin accumulations along axons are strongly abolished in the LatA treated example in B, whereas PMS are still visible. (**C**,**D**) Quantitative analysis of PMS abundance at different culture stages shows a gradual increase from 6HIV to 10DIV (C); PMS abundance normalised to parallel control cultures in neurons at 8HIV upon different pharmacological or genetic manipulations of actin and/or actin regulators as indicated (D); numbers in the bars are sample numbers (i.e. analysed regions of interest), error bars represent SEM of independent experimental repeats. Scale bar in A represents 1μm in A and B and 0.125μm in A’ to B”.

To summarise our results so far, PMS in young growing axons appears qualitatively the same, but still undergoes structural maturation, displays a higher dependence on polymerisation and becomes gradually more abundant over a period of days. Our data suggest that PMS (1) contains short actin filaments which are high in number and particularly reliant on nucleation processes, (2) requires adducin function likely to cap and stabilise the plus ends of actin filaments, and (3) needs spectrin for the formation of robust and abundant periodic patterns. These results provide experimental support for the cortical model for PMS that was originally proposed based on image data (Xu *et al.*, 2013). The consistency of our data with the original PMS model strongly suggests that the molecular structure of PMS is evolutionarily well conserved.

### F-actin has MT-maintaining roles in axons

We then started to explore the currently unknown, yet potentially important roles of PMS or the cortical actin they represent (from now on referred to as PMS/cortical axon) in axons.

**Figure 5.**
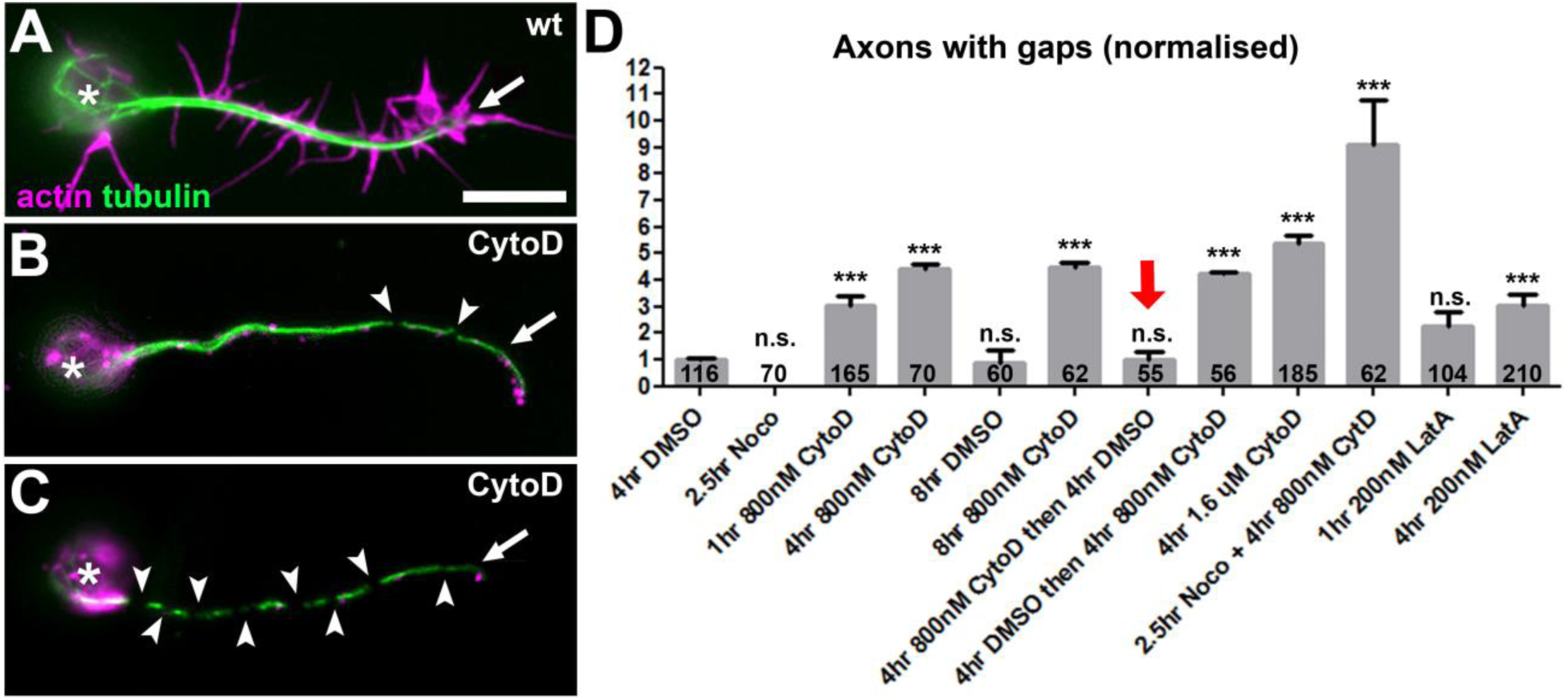
F-actin has MT stabilising roles. (**A-C**) Primary wildtype neurons at 8HIV stained with phalloidin for actin (magenta) and against tubulin (green), either untreated (wt) or treated with LatA or CytoD (asterisks, cell body; arrows, axon tips; arrowheads, MT gaps); images show gap phenotypes in C. (**D**) Quantification plotting axons with gaps normalised to 4hr DMSO-treated controls; note that washout of CytoD can revert the phenotype (red arrow; bar on its right shows that CytoD still has gap-inducing activity during the washout period) and that co-treatment with nocodazole (Noco) enhances the phenotype (highest bar); numbers in bars refer to analysed neurons; all data were compared to DMSO controls via X^2^ analysis (NS: P>0.05, *: P<0.05, **: P<0.01,***: P<0.001); for detailed data see Tab.S1. Scale bar in A represents 10μm in A-C.

Interestingly, we observed that, compared to controls, axons of neurons treated with the actin-destabilising drug CytoD showed frequent gaps in their MT arrays. There was a 4.5-fold increase in MT breaks or gaps upon 4hr treatment with 800nM CytoD (4-8HIV), and 9-fold increase when doubling the CytoD dose to 1.6μM (Fig. 5C,D). These gaps usually reflect unstable MTs and have previously been reported to occur under strong MT-destabilising conditions (Voelzmann *et al.*, 2016b). Importantly, these effects of CytoD were reverted when washing out the drug during a 4hr period (Fig. 5D), suggesting that F-actin mediates some sustained and acutely acting function in MT maintenance. In parallel, we observed that the CytoD-treated neurons showed a significant trend to lose axons. Thus, 29% of neurons (identified using the anti-ELAV antibody) tend to have no axon under normal conditions in our cultures. This number is only insignificantly increased to 31% upon 4hr treatment with 800nM CytoD, but to 45% of neurons when treating with 1.6μM CytoD for 4hrs (Fig. 6B,E).

This dose-dependent impairment of axonal MTs, which can culminate even in complete failure to form or maintain axons, suggests that F-actin has MT-protecting roles in axons. To test this notion further, we combined CytoD treatment with conditions that directly destabilise MTs and asked whether this would dramatically increase gaps and/or axon loss. As a baseline, we used neurons treated with 800nM CytoD for 4hrs, which showed only mild phenotypes compared to DMSO-treated neurons (4.5 times more axons with tubulin gaps; 31% of neurons without axons; see previous paragraph). When treating neurons first with 800nM CytoD for 4hrs (4-8HIV) and then co-treating with 20μM of the microtubule-destabilising drug nocodazole for the last 2.5hrs, the phenotypic defects almost doubled (5.5-8HIV; 9 times more axons with gaps; 51% of neurons without axon; Figs. 5D, 6E). In contrast, treating neurons with only nocodazole caused no obvious increases in tubulin gaps or axon loss (Figs. 5D, 6E) (*Sánchez-Soriano et al., 2009;* Alves-Silva *et al.*, 2012).

The effects of CytoD were similarly enhanced when affecting MT stability genetically, i.e. using the *shot*^*3*^ mutant allele to remove the MT-binding and -stabilising spectraplakin protein Short stop (Shot) (Yang *et al.*, 1999; Alves-Silva *et al.*, 2012). In agreement with earlier reports (Voelzmann *et al.*, 2016b), CytoD treatment has a strong effect on axonal MTs in these neurons: when treating *shot* mutant neurons for 4hrs with CytoD (4-8HIV at 800 nM), 64% of neurons lack axons, which is about double the amount observed in CytoD-treated wildtype (31%, see above) or in untreated *shot*^*3*^ mutant neurons (36%; Figs.6C,D,E). Therefore, *shot*^*3*^ mutant neurons, where MTs lack Shot-mediated protection, seem to benefit from parallel MT-maintaining functions of F-actin. In agreement with this conclusion, we found that *shot*^*3*^ mutant neurons display normal PMS abundance (Fig. 2F).

In conclusion, F-actin appears to protect axonal MTs through a sustained, acutely acting mechanism, and the power of this stabilising function becomes particularly apparent when weakening MT networks via nocodazole application or genetic removal of MT stabilising proteins such as Shot. Our results lead us to speculate that F-actin and Shot provide two independent, parallel mechanisms of MT stabilisation. Notably, 4hr LatA treatments (not affecting PMS at 10DIV; Fig. 2F) show milder MT gap and axon retraction phenotypes than CytoD treatments (strongly affecting PMS), making it tempting to speculate that the F-actin fractions contributing to MT maintenance are residing within PMS.

### Cortical F-actin acutely promotes MT polymerisation and acts in parallel to Shot

To understand how F-actin maintains MTs, we studied MT dynamics in axons upon drug application. For this we performed live imaging of EB1::GFP, which primarily localises as comets at plus ends of actively polymerising MTs (Fig.7A,B; suppl. movies S1 and S2) (Alves-Silva *et al.*, 2012).

**Figure 7.**
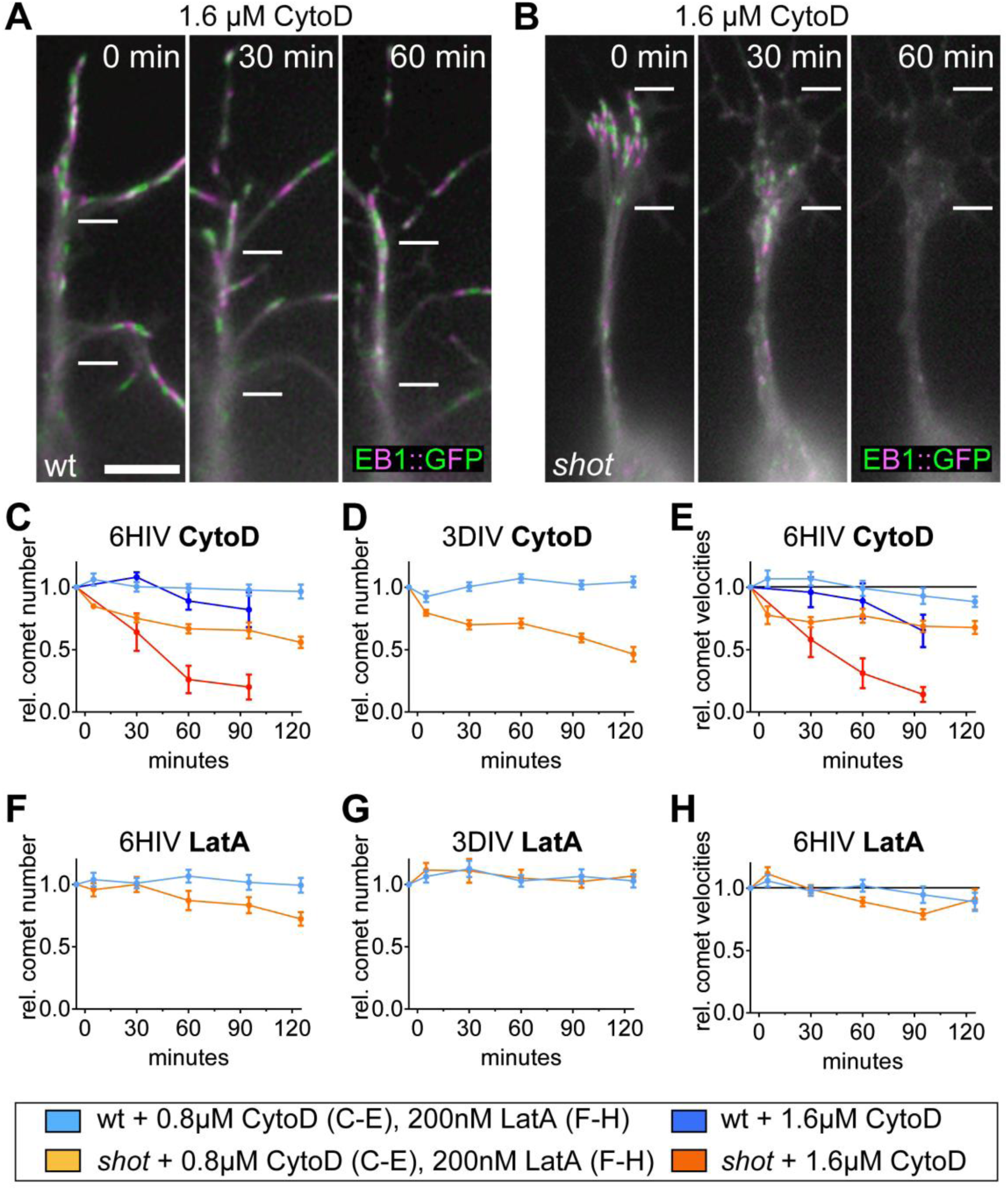
Live recordings of wildtype and *shot* deficient primary neurons expressing EB1::GFP. (**A**, **B**) Stills of movies of neurons at 6HIV taken at three time points (0, 30 and 60 min after treatment), where each still is a projection of four images which are 3s apart and alternately coloured in green and magenta to indicate the movement of EB1::GFP comets; note that comets come to a halt and vanish only in *shot* mutant neurons. Measurements of comet numbers (**C**, **D**, **F**, **G**) and velocities (**E**, **H**) of EB1::GFP comets in wild-type and *shot*^*3*^ mutant neurons upon treatment with either 0.8μM CytoD, 1.6μM CytoD or 200nM LatA respectively, as indicated in box below; velocity of WT is 0.154im/s±0.01SEM; for detailed data see Tab.S1. Scale bar indicates 5μm.

We first treated wildtype neurons at 6HIV or 3DIV with 800nM or 1.6μM CytoD and measured the direct impact on the numbers and velocities of EB1::GFP comets (blue curves in Fig. 7C-E). Comet numbers were affected little or not at all by CytoD treatment at both culture stages (Fig. 7C,D), but at 6HIV the velocity of EB1 comets gradually decreased, and this effect was clearly dose dependent (Fig. 7E). This decrease is likely to reflect reduced net polymerisation of MTs and could explain the dose-dependent gaps observed via axonal tubulin staining in CytoD-treated wildtype neurons at that stage (Figs. 3B,5D and 7C,E). Similarly, application of LatA to wildtype neurons (blue lines in Fig.7 F-H) had no effect on comet numbers initially, however after one hour we observed a mild drop in comet velocity, in agreement with our observations that 1hr LatA treatment of young neurons has little impact on PMS abundance and axonal gaps, but that effects are observed after 4hr treatment (Figs.4D, 5D). Therefore, acute removal of PMS correlates with an acute reduction in MT polymerisation.

These data for wildtype neurons reveal fairly mild effects. In contrast, the responses become very pronounced when repeating the same experiments in *shot*^*3*^ mutant neurons (orange lines in Fig. 7). Thus, upon application of CytoD to *shot*^*3*^ mutant neurons at either 6HIV or 3DIV, comets instantaneously changed from steady propagation to stalling behaviour and then gradually faded away; these effects were enhanced when increasing CytoD concentration from 800nM to 1μM (Fig. 7C-E; Suppl. Movie S2. Detailed image analyses revealed a significant loss of comet numbers and significant slow-down of comet speed in *shot*^*3*^ mutant neurons at all stages analysed (Fig. 7C-E). In contrast, application of LatA to *shot*^*3*^ mutant neurons had no effect on comet number and velocity at 3DIV consistent with the fact that PMS are not affected under these conditions (Fig. 7G). At 6HIV, there seems to be a delayed drop in comet numbers and velocity in *shot*^*3*^ mutant neurons (Fig. 7F,H), which would be consistent with our observation of a delayed decrease in PMS abundance upon LatA application (Figs. 2F,4D).

These data further suggest a role for PMS/cortical actin in MT maintenance and the existence of a sustained mechanism downstream of PMS/cortical actin which acts through stabilising MT polymerisation or preventing its inhibition. The data also support our notion that Shot provides a parallel stabilising mechanism that likewise sustains MT polymerisation; and both mechanisms seem to be able to compensate for each other.

### Reduced PMS abundance correlates with changes in axonal MT organisation

To analyse relations between PMS and MTs in a quantifiable manner, we used a further phenotype of *shot*^*3*^ mutant neurons, consisting in varicose regions of axons which are formed by disorganised MTs that are not arranged into bundles but curl up and criss-cross each other (Fig. 8A) (Sánchez-Soriano *et al.*, 2009; Voelzmann *et al.*, 2016a). Measurements of these areas are easy to perform and statistically robust across large neuron populations and we express them as “MT disorganisation index” (MDI; see Methods). The MDI is measured most effectively at 6-8HIV, still reliably in mature neurons up to about 3DIV, and increasingly unreliable at 10DIV when the total length of axons becomes difficult to determine within the densely grown axon networks (Fig. 1E,F).

**Figure 8.**
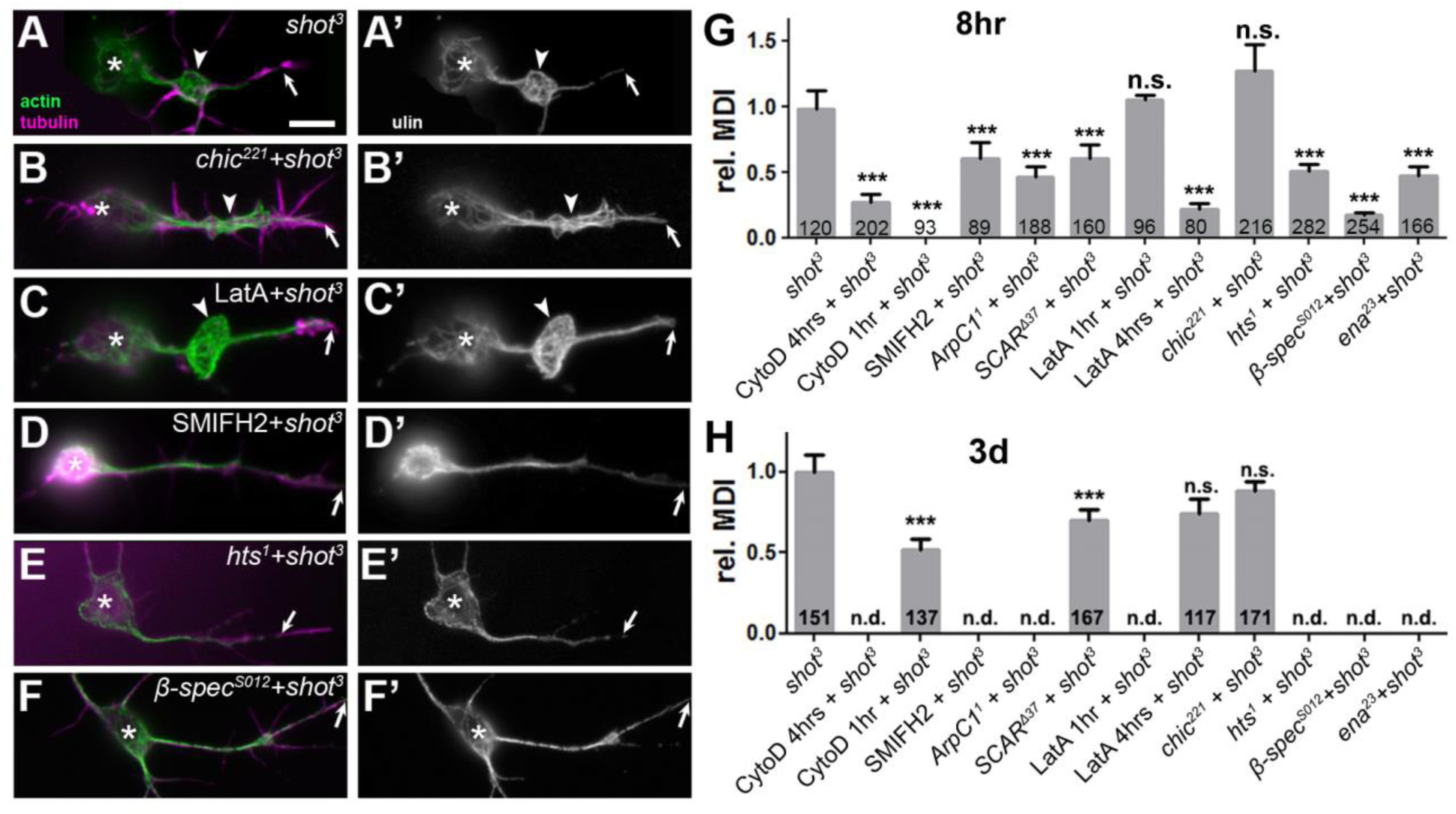
F-actin manipulations in *shot*^*3*^ mutant neurons. (**-F**’) *shot*^*3*^ mutant primary neurons at 8HIV stained for tubulin (green) and actin (magenta) combined with different actin manipulation as indicated (asterisks, cell bodies; arrows, axon tips; arrowheads, areas of MT disorganisation). (**G**,**H**) Quantifications of MDI for neurons at 8HIV (H) and 3DIV (I); numbers in bars refer to neurons analysed; all data normalised to *shot;* for detailed data see Tab.S1. P values were calculated using the Mann-Whitney Rank Sum test (NS: P>0.05, *: P<0.05, **: P<0.01, ***: P<0.001). Scale bar in A represents 10μm in A-G’.

Using CytoD, we tested whether removal of PMS/cortical actin in *shot*^*3*^ mutant neurons affects the MDI, and we predicted a reduction due to the observed reduction in MT polymerisation (Fig. 7). When treated with 800nM CytoD from 4 to 8HIV, those neurons which retained an axon no longer showed disorganisation (MDI = 0; all compared to untreated *shot*^*3*^ mutant neurons; Fig. 8G), and axons often appeared very thin (Fig. 6D). This means that either MTs in these axons become reorganised into bundles or they vanish, of which the latter would be consistent with the overall loss of axons (Fig. 6). At 3DIV, 4hr treatments with 800nM CytoD significantly reduced the MDI by 48% (Fig.8H). This phenotype is milder than at 8HIV and might be explained by the fact that MT polymerisation (the process most likely affected by combined loss of Shot and F-actin; Fig. 7) is less frequent in old neurons than young ones (8HIV: 0.68±0.05 comets/μm axon length; 3DIV: 0.29±0.04; n=20, pMW<0.0001). Therefore, inhibiting MT polymerisation at 3DIV would be expected to show a slower impact on varicose areas.

Also LatA treatment of *shot^3^* mutant neurons revealed a correlation between MDI and PMS abundance (Figs. 2 and 4): at 3DIV, no significant MDI reduction was observed upon 4hr treatment with 200nM LatA (Fig. 8H, 2F); at 8HIV, a preceding 1hr LatA treatment had no effect but a 4hr treatment reduced MDI to 22% (Figs. 8C,G and 4D). These MDI results at 8HIV are paralleled by our observations in LatA-treated wildtype control neurons, where MT gaps and axon loss were not very obvious after 1hr but became prominent after 4hr LatA treatment (Figs. 5D and 6E), also suggesting that a reduction in MDI might involve loss of MTs.

We also found a correlation between MDI and PMS abundance when analysing neurons in which the *shot*^*3*^ mutant allele was combined with various mutations of actin regulators. For example, the *shot*^*3*^ *SCAR*^Ä37^ double-mutant neurons showed a highly significant reduction in MDI at both 8HIV and 3DIV, correlating well with the significantly reduced PMS abundance observed in *SCAR*^Ä37^ mutant neurons. In contrast, *shot*^*3*^ *chic*^*221*^ double-mutant neurons showed no significant MDI reduction (Fig. 8A,G,H), in agreement with the fact that the *chic*^221^ mutation does not affect PMS abundance (Fig. 2). Therefore, these results also correlated well.

In total we analysed the MDI in 16 different conditions where drugs or actin-regulator mutations were combined with *shot*^*3*^ mutant background (Fig. 8). We then plotted these MDI data (Fig. 8) against the respective PMS abundance data obtained for wildtype neurons at 8HIV and 10DIV (Figs. 2 and 4), and found a highly significant correlation between the presence/absence of PMS and high/low values for MDI (Figs. 9B; Spearman correlation coefficient = 0.782, p = 0.0009; Tab. S1). These correlations suggest that MT-regulating capacity is a function of the amount of PMS present in axons. Associating PMS with this function becomes even more convincing when considering that filopodial readouts (representing long actin filament networks) are not at all correlated with MT phenotypes.

**Figure 9.**
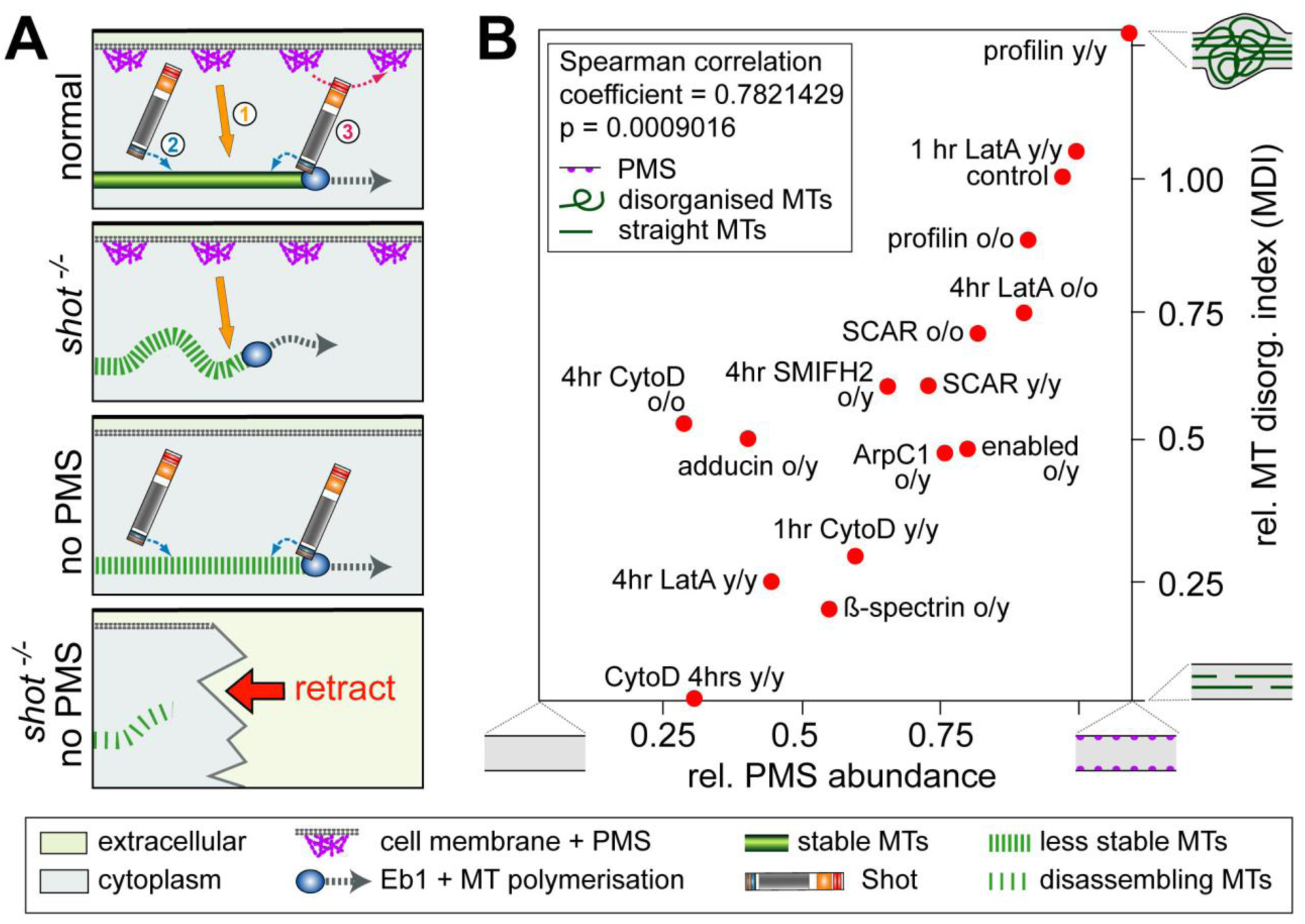
Presence of cortical actin correlates with MDI values. (**A**) Schematics illustrating roles of actin in MT stabilisation (1; orange arrow; this paper) and two independent roles of Shot in maintaining coalescent MT bundles (Alves-Silva *et al.*, 2012): through MT/MT bundle stabilisation (2; blue arrow) and guidance of polymerising MTs into parallel bundles (3). Upon Shot deficiency MTs become less stable and disorganised, upon PMS loss MTs become less stable, loss of both leads to MT disassembly and axon retraction (symbols explained in box below). (**B**) Correlation plot comparing degrees of PMS abundance (data from Figure 2) with degrees of MDI (data from Figure 8); all conditions are in combination with loss of Shot function (*shot^3^* mutant allele in homozygosis); in each pairing “o” indicates old neurons (3 or 10DIV) and “y” young neurons (6-8HIV) with regard to PMS analysis (before slash) and MDI analysis (after slash); extreme phenotypes for PMS abundance and MDI are symbolised as explained in inset.

## Discussion

### Newly discovered roles for PMS/cortical actin in axonal MT maintenance

Axonal MT bundles provide the essential structural backbones and highways for life-sustaining transport in axons, and damage to these bundles seems to correlate with axonal decay (Adalbert and Coleman, 2012). Active maintenance including constant polymerisation and disassembly of MTs is required to prevent decay through wear and tear, but hardly anything is known about these mechanisms (Voelzmann *et al.*, 2016a). Here we report that axonal actin and the spectraplakin protein Shot have independent, complementary roles in regulating and maintaining MT polymerisation in growing and mature axons, and we show that the combined absence of these mechanisms induces a severe axon loss. Our data strongly suggest that the actin fraction responsible for this regulation is provided by PMS, because only in conditions where PMS are affected do we observe MT defects, including gaps in tubulin staining, loss of MTs accompanied by axon retraction, inhibition of MT polymerisation and a reduction in disorganised MTs (measured as MDI; Fig. S4).

Of these readouts, the MDI data were assessed quantitatively and revealed a tight correlation with PMS abundance, suggesting that MT regulatory mechanisms act as a function of the amount of PMS/cortical actin present in axons. We suggest that the process underlying MDI reduction is MT loss rather than MT reorganisation. First, the size of areas with MT disorganisation (expressed as MDI) tightly correlates with EB1::GFP comet numbers (see Figs.7 and S3). Since each comet represents a single MT, the size of varicose areas seems to be a function of MT number and content. Second, severe loss of MDI occurs together with loss of whole axons. Third, we know that combined loss of PMS and Shot causes loss of MT polymerisation, providing a likely mechanism leading to MT gaps, loss of axons and reduction in MDI. Unfortunately, direct measurements of MT numbers in axons are impossible at the light microscopic level (Mikhaylova *et al.*, 2015) and will require high-pressure freezing EM as the currently best method to provide high resolution combined with good preservation of all cell structures including unstable MTs (McDonald, 2007).

Axon maintenance requires continued MT polymerisation and disassembly to prevent senescence of MT bundles in axons (Voelzmann *et al.*, 2016a). In this scenario, the roles of PMS/cortical actin in sustaining MT polymerisation proposed here, offer conceptually new potential explanations for brain disorders linked to gene mutations of cortical actin regulators: adducin (ADD3) is linked to human cerebral palsy (Kruer *et al.*, 2013), ß-spectrin to spinocerebellar ataxia (SPTBN2; OMIM ID: 600224, 615386), ankyrin to mental retardation (ANK3; OMIM ID: 615493), actin itself to Baraitser-Winter syndrome and dystonia with neurodegenerative traits (ACTB7; OMIM ID: 243310, 607371). Any of these pathologies could potentially relate to gradual MT bundle decay in axons through a reduction in MT polymerisation and turn-over. In support of our interpretation, Spectrin deficient neurons were shown to display axon breakage in *C. elegans* (Hammarlund *et al.*, 2007), cause axonal transport defects coupled to neurodegeneration in *Drosophila* (Lorenzo *et al.*, 2010), and loss of *Drosophila* adducin or spectrin causes synapse retraction *in vivo* (Pielage *et al.*, 2005; Pielage *et al.*, 2011). Likewise, spectraplakins as the second important promoters of MT polymerisation link to the neurodegenerative disorder type IV hereditary sensory and autonomic neuropathy (OMIM ID: 614653). Whilst a number of mechanisms were proposed (Alves-Silva *et al.*, 2012; Ferrier *et al.*, 2013; Voelzmann *et al.*, 2016a), weakening of the MT polymerisation machinery might be a further mechanism underlying this disorder.

Interestingly, the same manipulations as used by us (i.e. *shot* mutant conditions in flies, CytoD treatment in mammalian neurons) activate the kinase DLK which, in turn, promotes axon injury responses including axon degeneration (Valakh *et al.*, 2013). DLK might therefore be involved in axonal retraction observed in this work, for example in response to MT stress caused by impaired MT polymerisation.

Our finding that co-depletion of F-actin and Shot causes axon loss also provides interesting new ideas for axon stump degeneration after injury. This event has been associated with an elevation of intracellular free calcium levels leading to calpain-mediated removal of actin-spectrin (Bradke *et al.*, 2012). Notably, the same calcium events can be expected to trigger detachment of Shot/spectraplakins from MTs through binding to their C-terminal EF-hand motifs (Wu *et al.*, 2011; Kapur *et al.*, 2012; Ka *et al.*, 2014). Therefore, calcium can synchronously remove two essential MT-stabilising mechanisms and should therefore cause axon retraction, as observed in our experiments.

### Axonal PMS as a promising cell model for the study of cortical actin in neurons

The strategy we chose was to use SIM imaging which provides slightly less resolution than STORM or STED but is fast and reliable, and therefore ideally suited for quantitative analysis with very high sample numbers. This enabled us to apply a wide range of genetic and pharmacological manipulations and classify their affects using PMS abundance as readout. PMS abundance is a functionally relevant parameter as can be deduced from its close correlation with parallel effects on MT bundles, MT dynamics and even axon maintenance. However, it remains to be seen whether the regular repetitive pattern *per se* is relevant for MT maintenance, or rather the mere presence of cortical F-actin. For example, the periodic arrangement of PMS could either be the outcome of active developmental mechanisms or the mere consequence of tubular axon morphology. Axons are cylinders that display longitudinal contraction (Bray, 1984; Siechen *et al.*, 2009), potentially providing conditions under which diffusely cross-linked cortical actin networks could shuffle into linear, periodic patterns - as was similarly proposed for actin rings in tracheal tubes (Hannezo *et al.*, 2015). If tubular structure is a key prerequisite for periodic actin arrangements, this would explain why dendritic and even neurite-like glial processes have actin rings.

Even if we still have to unravel the processes through which PMS are assembled, the mere existence of PMS provides us with powerful readouts for studying the regulation and function of cortical actin in neurons.

### Future prospects

Here we propose an important role for PMS/cortical actin in MT maintenance with potential implications for neurodegenerative diseases linking to cortical actin factors. The challenge now will be to find the mechanisms linking cortical actin networks to MT polymerisation. This will not be a trivial task when considering that MT polymerisation is coordinated by a complex machinery which involves the regulation of tubulin supply, a large number of proteins associating with MT plus ends, as well as shaft-based mechanisms (Voelzmann *et al.*, 2016a).

Apart from roles in MT maintenance, PMS/cortical actin might contribute to axon biology through further relevant mechanisms: (1) to guide polymerising MTs into ordered parallel bundles, mediated by the spectraplakin actin-MT linkers (Sánchez-Soriano *et al.*, 2009; Alves-Silva *et al.*, 2012; Prokop *et al.*, 2013), (2) to anchor the minus ends of MTs (Nashchekin *et al.*, 2016; Ning *et al.*, 2016), (3) to serve as anchor for Dynein/Dynactin-mediated sliding of MTs and transport of MT fragments (Myers *et al.*, 2006), (4) to anchor and compartmentalise transmembrane proteins along axons (relevant for action potentials or the adhesion to ensheathing glia) (Baines, 2010; Machnicka *et al.*, 2014; Albrecht *et al.*, 2016; Zhang *et al.*, 2016), and (5) perhaps even to contribute to the regulation of collateral branching (Kalil and Dent, 2014). All these functions can now be studied using PMS as powerful readouts and building on available concepts for the structure and function of cortical actin derived from this work and from studies in mammalian axons (see Introduction) or from non-neuronal systems such as erythrocytes (Baines, 2010).

## Acknowledgement

This work was made possible through funding by the BBSRC (BB/L000717/1, BB/M007553/1) to A. P, as well as support by parents and Manchester’s Faculty of Life Sciences (now FBMH) to Y.Q, and a Leverhulme Early Career Fellowship to S.P.P. Microscopes at the Bioimaging Facility in Manchester were purchased with grants from BBSRC, The Wellcome Trust and the University of Manchester Strategic Fund, and the Fly Facility has been supported by funds from The University of Manchester and the Wellcome Trust (087742/Z/08/Z). Structured illumination and STED microscopes at the Research Complex at Harwell were funded by the MRC (MR/K015591/1) and BBSRC (BB/L014327/1) and imaging time was made possible through three successive grants by the STFC to A.P. We thank Christopher Tynan for his support with STED imaging, Andre Voelzmann, Natalia Sánchez-Soriano and Tom Millard for helpful comments on the manuscript, and many colleagues and the Bloomington *Drosophila* Stock Center (NIH P40OD018537) for kindly providing stocks and materials.

